# Measuring kinetics and metastatic propensity of CTCs by blood exchange between mice

**DOI:** 10.1101/2020.11.01.350918

**Authors:** Bashar Hamza, Alex B. Miller, Lara Meier, Max Stockslager, Emily M. King, Sheng Rong Ng, Kelsey L. DeGouveia, Nolawit Mulugeta, Nicholas L. Calistri, Haley Strouf, Lin Lin, Christopher R. Chin, Robert A. Weinberg, Alex K. Shalek, Tyler Jacks, Scott Manalis

**Author notes:** co-first author (Bashar Hamza, Alex B. Miller).

## Abstract

Existing pre-clinical methods for acquiring dissemination kinetics of rare circulating tumor cells (CTCs) en route to forming metastases have not been capable of providing a direct measure of CTC intravasation rate and subsequent half-life in the circulation. Here, we demonstrate an approach for measuring endogenous CTC kinetics by continuously exchanging CTC-containing blood over several hours between un-anesthetized, tumor-bearing mice and healthy, tumor-free counterparts. By tracking CTC transfer rates using an autochthonous small cell lung cancer model, we extrapolated half-life times in the circulation of 50-100 seconds and intravasation rates between 4,000 and 27,000 CTCs/hour – an average daily shedding rate equivalent to ∼0.07% of the total number of primary tumor cells in the lung. Additionally, transfer of 1-2% of daily-shed CTCs from late-stage tumor-bearing mice generated macrometastases in healthy recipient mice. We envision that our technique will help further elucidate the role of CTCs and the rate-limiting steps in metastasis.

## Introduction

Circulating tumor cells (CTCs) **—** cells shed into the bloodstream from primary and metastatic tumor deposits **—** represent the intermediary component of the metastatic cascade. Measuring their intravasation rate and half-life time in the circulatory system, which together govern their blood levels, has been an important step towards elucidating the kinetics of their seeding of distant tissues and subsequent outgrowth of metastatic colonies. Traditionally, the fate of tumor cells has been examined by injecting tumor cell lines intravenously into animal models (mostly mice, rats, and rabbits)^1–4^. By analyzing terminal blood and other major organs from multiple animals at different time points post inoculation, these initial studies suggested extremely short half-life times in circulation (less than 1 second^3,5^). The intravenously injected cells seemed to arrest in the capillaries of the first organs they encountered almost immediately after injection, but their subsequent proliferation into secondary lesions was influenced by host-tumor cell interactions operating within specific organs. Although these studies established the basis of the “seed and soil” hypothesis^6–8^, these methods have not been amenable to endogenously generated CTCs that originate from solid primary tumors, which would be expected to exhibit different physiology than CTCs derived from established cell lines. Additionally, previous methods^9–19^ have not yet provided direct measures of the intravasation rate for CTCs.

Beyond the circulatory dynamics, a quantitative functional assessment of CTCs’ intrinsic propensity to proliferate in the parenchyma of distant organs is important for identifying the biological properties of metastasis-initiating cells. Current pre-clinical attempts to address this aspect rely on either murine cell lines or CTCs isolated from patient blood samples *ex vivo* prior to their injection into immunocompromised animals^20–25^. While these various studies presented different approaches for growing tumors in laboratory animals to study metastasis or explore different potential therapeutic options, newer pre-clinical methods utilizing immunocompetent mice and requiring less *ex vivo* manipulation of CTCs are likely to facilitate a deeper understanding of the role of CTCs in metastasis.

To address these limitations, we developed a novel blood-exchange method between pairs of un-anesthetized mice for studying the circulatory dynamics and tumorigenicity of CTCs. We applied our method to an autochthonous, genetically engineered mouse model (GEMM) of small cell lung cancer (SCLC) to study the kinetics of endogenous CTC generated from tumors that arise *de novo* in the context of surrounding stroma and a fully functional host immune system^26^. Unlike parabiosis, our method does not require a permanent surgical connection between the vasculatures of the two mice. Instead, blood is temporarily exchanged for several hours through the implanted catheters. The blood-exchange method presented here will enable a series of experiments that can answer fundamental questions about the relationship between CTC characteristics and metastasis.

## Results

### Blood Exchange for Direct CTC Kinetics Studies

To measure the CTC half-life time and generation (intravasation) rate, we utilized a syngeneic non-tumor-bearing “healthy” mouse (HM) as a recipient for CTCs generated by a tumor-bearing mouse (TBM). In this experimental model, primary tumors in the TBM are initiated by deletion of the *Trp53, Rb1*, and *Pten* (PRPten) tumor suppressor genes in the murine lung epithelium^27^ and include a Cre-activated tdTomato allele that engenders fluorescence in all tumor cells after tumor initiation, including derived CTCs. TBMs survive for approximately 6 months post tumor initiation with detectable primary (lung) and metastatic (liver) tumors starting at approximately 5 months post tumor initiation.

In our experimental setup (**Fig. 1a**), CTC-containing blood is exchanged between a TBM (5-6 months post tumor initiation) and an HM counterpart of the same sex and similar age and weight. The exchanged blood is monitored in real time through two CTC Counters (see *Methods*), being passed at a flow rate of 60 µL/min in a closed-loop manner (**Supplementary Figs. 1 and 2**). Within each CTC Counter, fluorescent CTCs are excited using a 532 nm dual*-*excitation laser beam configuration focused across the microfluidic blood-flow channel near the inlet of the device (**Supplementary Fig. 1** and **3**). The resulting dual emission peaks from the photomultiplier tube are recorded for offline peak analysis and validation purposes (**Fig. 1b**).

**Fig. 1.**
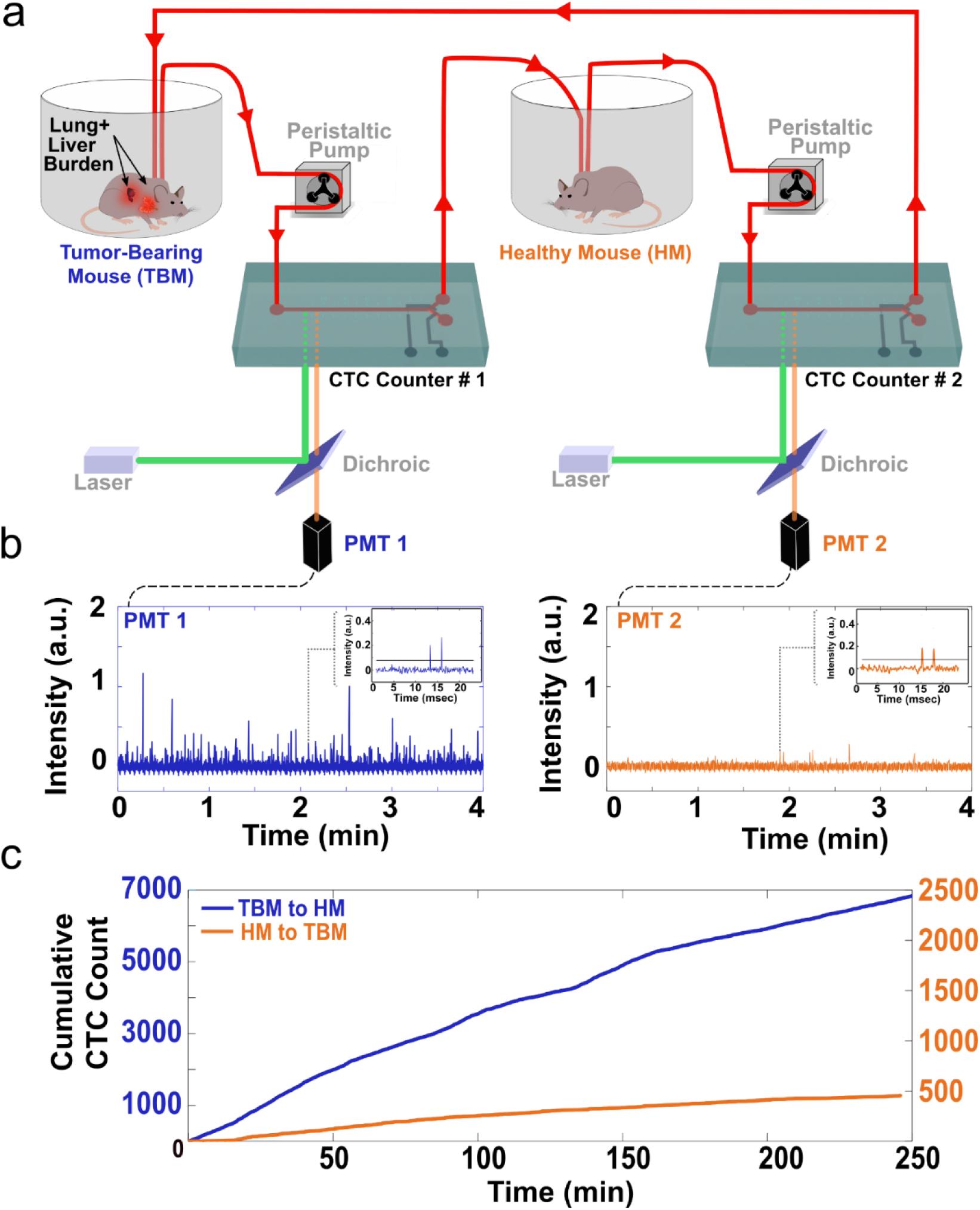
A blood-exchange method for direct CTC kinetics studies. (**a**) Schematic demonstrating the blood exchange method in which the circulatory systems of two mice (one tumor bearing mouse (TBM) and one healthy mouse (HM)) are connected in a closed loop through two CTC counters. Two peristaltic pumps set to identical flow rates push the blood around the system and through the CTC counters. For each CTC counter, a series of 2 laser lines is used to excite the flowing genetically-fluorescent CTCs (See Supplementary Fig. 1-3 for more details). Emitted signal is directed through a dichroic filter toward a photomultiplier tube (PMT) for detection. (**b**) PMT lowpass-filtered spectra demonstrate a series of detected CTCs by each of the two CTC counters (PMT 1 = CTC counter 1 (blue), PMT 2 = CTC counter 2 (orange)). Inset demonstrates the dual peak configuration created when each CTC passes under the two laser lines projected across the flow channel. (**c**) Cumulative CTC counts over time from the two CTC counters demonstrating the higher injection rate (blue trajectory with left Y axis) compared to the return rate (orange trajectory with the right Y axis).

**Fig. 2.**
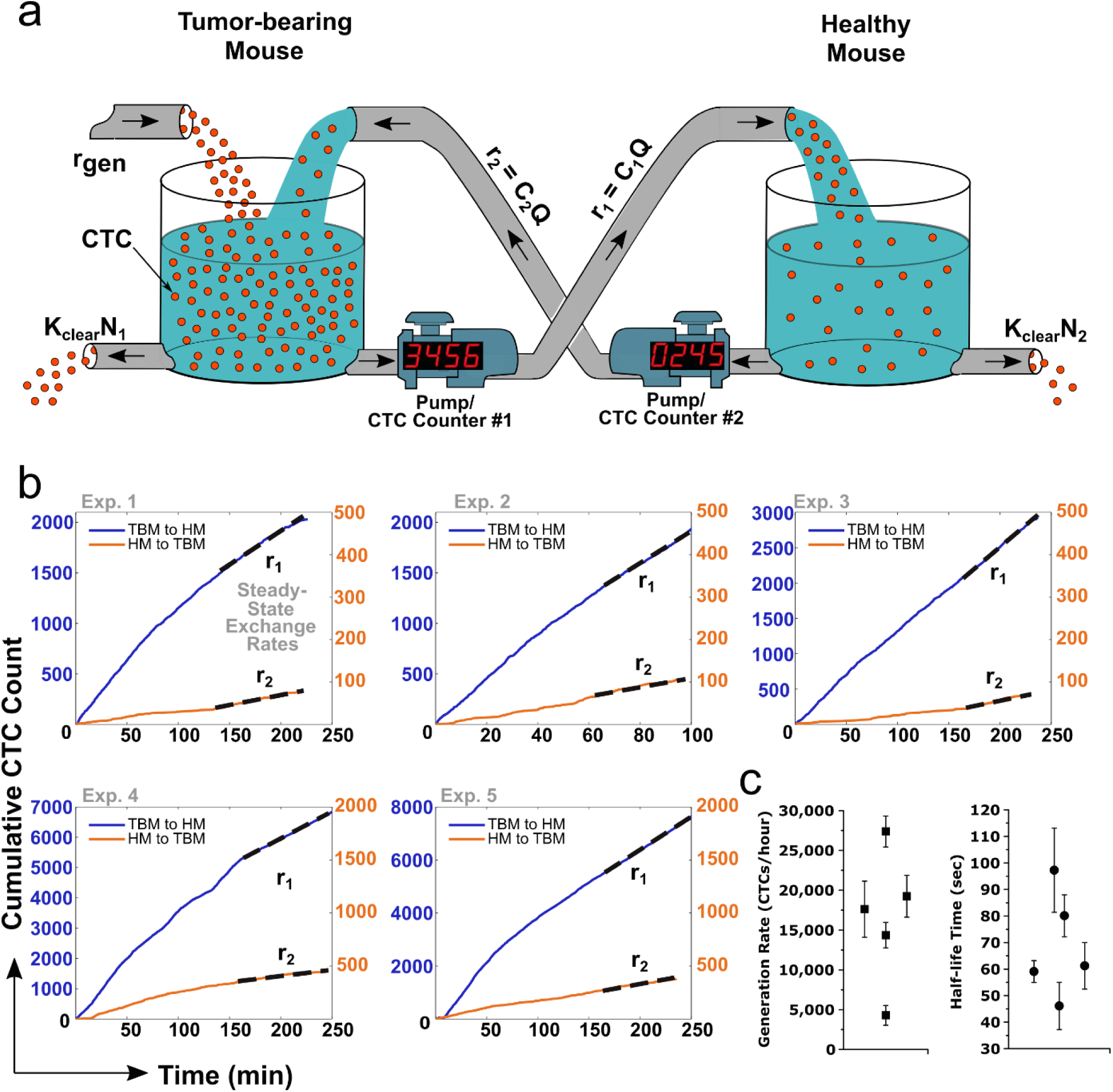
An analytical model for extracting CTC generation rate and half-life time in blood. (**a**) A visual representation of the relevant parameters of the blood exchange technique to solve for the generation rate and the half-life time of CTCs. The circulatory system of each mouse is represented as a well-mixed container of red spheres (CTCs). In the TBM (left tank) CTCs enter the circulation from the tumor microenvironment at a rate equal to ***r***_***gen***_. CTC clearance out of the circulation is represented by a hole at the bottom of each container with a clearance rate of ***K***_***clear***_ ***× n***. Pumps with counters represent the CTC counter systems and their peristaltic pumps that transfer the CTC-containing blood at rates equal to ***C × Q***. (**b**) Five cumulative CTC count charts over time representing the counted CTCs when exchanged between five different pairs of mice (5 late-stage TBMs and 5 HMs). In each experiment a TBM is connected to an HM of the same sex, age, and genotype. Dotted lines represent the best fits to extract the steady-state exchanged rates between the two mice (***r***_***1***_ and ***r***_***2***_). (**c**) Scatter plots for the calculated CTC generation rates and the half-life times of each of the five blood-exchange experiments. Error bars represent the propagated error due to the absolute uncertainty (s.d.) in the ***r***_***1***_ and ***r***_***2***_ estimates.

To access the circulation, mice undergo a cannulation surgery to externalize two catheters from the left carotid artery and the right jugular vein. Continuous blood flow between the two un-anesthetized mice is controlled by two peristaltic pumps and can be executed over several hours until steady-state CTC exchange rates are measured. The relatively high heart rate in mice (400-600 beats per minute resulting with high cardiac outputs of ∼20 mL/min^28^) and low total circulating blood volume (∼1.5-2 mL) allow us to assume that CTCs are uniformly distributed in the bloodstream of each mouse when sampled continuously at the much lower volumetric flow rate of 60 µL/min between the two mice. This assumption was validated empirically by observing similar CTC concentrations in flowing blood during real-time scans and terminal blood samples collected by cardiac puncture from four SCLC-bearing animals (**Supplementary Table 1**).

In more detail, CTCs drawn from the TBM’s carotid artery line pass through the first, in-line CTC counter (labeled as CTC counter #1 in **Fig. 1a, Supplementary Fig. 2**) for real-time enumeration prior to their introduction into the jugular vein catheter of the HM. The total transport time of individual CTCs within the tubing from one circulatory system to the other is approximately 2 minutes at the 60 µL/min flow rate. CTCs that remain in the circulation of the HM are detected and counted by the second CTC counter (#2 in **Fig. 1a, Supplementary Fig. 2**) prior to their return to the right jugular vein of TBM. Once the raw PMT data are processed for validation of CTC counts (**Fig. 1b**), the cumulative counts over time from each CTC counter are then plotted to extract the CTC exchange rates necessary for calculation of the half-life time and the generation rate of CTCs by the primary tumor. **Fig. 1c** shows an example of one of these blood-exchange experiments, in which the HM received approximately 7,000 CTCs over the course of 4 hours and returned approximately 500 CTCs to the TBM during this time period.

### Modeling the Blood-Exchange Kinetics

Analysis of the numbers of CTCs exchanged in each direction between the tumor-bearing and healthy mice allows us to monitor the instantaneous concentration of CTCs in each mouse, which in turn enables us to estimate the rate at which CTCs are being shed from the tumor and cleared thereafter from the circulation. To estimate the CTC generation rate and half-life time, we fit our data to a simple model in which CTCs (represented by the red spheres in **Fig. 2a**) are generated from the lung tumor microenvironment at a constant rate (*r*_*gen*_) in the TBM and are cleared from the circulation of both mice by either lodging into capillaries across the different organs or by elimination through cell death (represented by a hole at the bottom of the tanks in **Fig. 2a**).

We assume first-order kinetics where CTCs remain in circulation for a half-life time of *t*_1/2_, or equivalently, are cleared with rate constant *K*_*clear*_ = ln(2) /*t*_1/2_. The two coupled differential equations describing how the number of CTCs in each mouse change with time are:

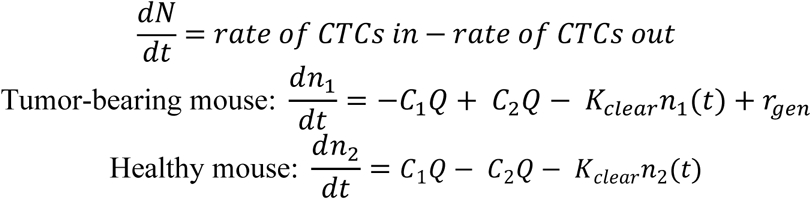

where *Q* is the constant volumetric flow rate of the pump exchanging the blood of the mice, *V* is the total blood volume of each mouse, and *c*_1_(*t*) = *n*_1_(*t*)/*V* and *c*_2_(*t*) = *n*_2_(*t*)/*V* are the concentrations of CTCs in the tumor-bearing and healthy mice, respectively.

Each CTC counter (represented by the pump with a digital counter in **Fig. 2a**) measures the *rate* at which CTCs pass from one mouse to another, which we assume to be proportional to the concentration of CTCs in the blood: *r*_1_(*t*) = *c*_1_(*t*)*Q* and *r*_2_(*t*) = *c*_2_(*t*)*Q* for the tumor-bearing-to-healthy CTC counter #1 and the healthy-to-tumor-bearing CTC counter #2, respectively. As discussed above, this “well-mixed” circulatory system assumption is justified, since the cardiac output in a small total blood volume is approximately 50-fold greater than the blood exchange flow rate between the mice of 60 μL/min.

Given measurements of *r*_1_and *r*_2_, the steady-state rates at which CTCs pass through both CTC counters, we can estimate the CTC generation rate and half-life in the circulation as:

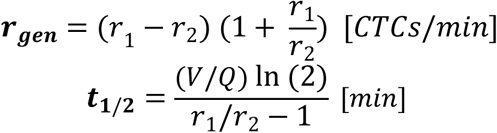

To validate this model, we performed five blood-exchange experiments between five different pairs of tumor-bearing mice and healthy counterparts (**Fig. 2b**). At the beginning of each experiment, we performed an initial scan (for at least 30 minutes) in which each mouse’s blood was scanned separately to ensure proper CTC counter functionality, reliable CTC detection, and stable blood flow from the carotid artery through the CTC counter and back into the jugular vein. Afterwards, the blood-exchange process was executed for at least two hours to allow for a sufficient time period for the CTC exchange rates to stabilize. During these five blood exchange experiments, the total number of CTCs transferred from the TBM to the HM varied from ∼2,000 to ∼8,000 CTCs (blue trajectories in **Fig. 2b**); 3-7% of these CTCs were returned from the HM to the TBM (orange trajectories in **Fig. 2b**). In order to extract the steady-state exchange rates from the empirical data, a best fit line (black dotted line in **Fig. 2b**) was applied to the cumulative count trajectories during the final interval of the blood-exchange experiments. During this interval, changes in CTC counts over time in each mouse (*dn*/*dt*) were approximately zero (i.e., ***r***_***1***_ and ***r***_***2***_ are roughly constant). CTC exchange rates (***r***_***1***_ and ***r***_***2***_) were assessed from different steady-state intervals (30, 45, 60, 75, and 90 minutes) to extrapolate the average and uncertainty (error) values for ***r***_***gen***_ and ***t***_***1/2***_ parameters (**Fig. 2c**; *Methods*). These ***r***_***1***_ and ***r***_***2***_ estimates are shown in **Supplementary Fig. 4a**.

**Fig. 3.**
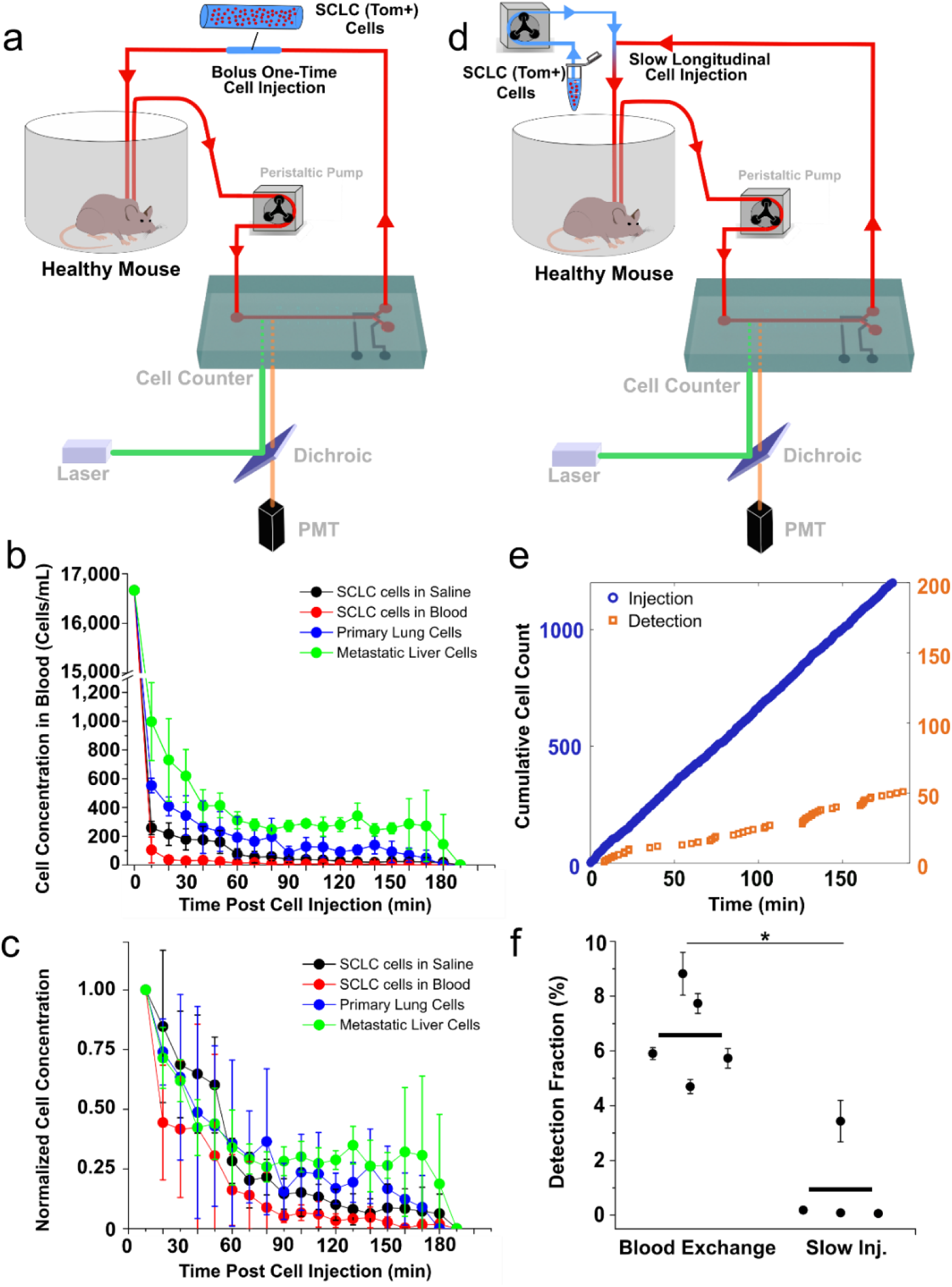
Comparison of CTC circulatory kinetics to SCLC cell line. (**a**) Schematic demonstrating the bolus cell-injection experiment configuration. A short tubing filled with ∼40 µL of blood or saline containing 25,000 cells of either SCLC cell line, dissociated primary tumor cells, or dissociated liver metastatic cells is added to the blood return line at the beginning of the experiment for a direct injection of its contents into the circulation within ∼1 minute. (**b-c**) Clearance kinetics plots representing the real-time concentration (**b**) and the normalized concentration to the initial (first 10 minutes) detected concentration. (**d**) Schematic demonstrating the slow cell-line injection experiment configuration. A second peristaltic pump is used to slowly infuse cell-containing saline into the circulation at a flow rate of 2-3 µL/min, with total injected cells mimicking that of the CTC blood exchange experiments. (**e**) Cumulative injection (blue) and detection (orange) cell counts over time. (**f**) Detection fraction plot representing the average fraction of total detected cells to the total injected cells in the last 30, 45, and 60 minutes of five blood-exchange experiments and four slow-injection experiments. All values are represented as mean ± sd. *p<0.05 (Mann-Whitney-Wilcoxon non-parametric test).

**Fig. 4.**
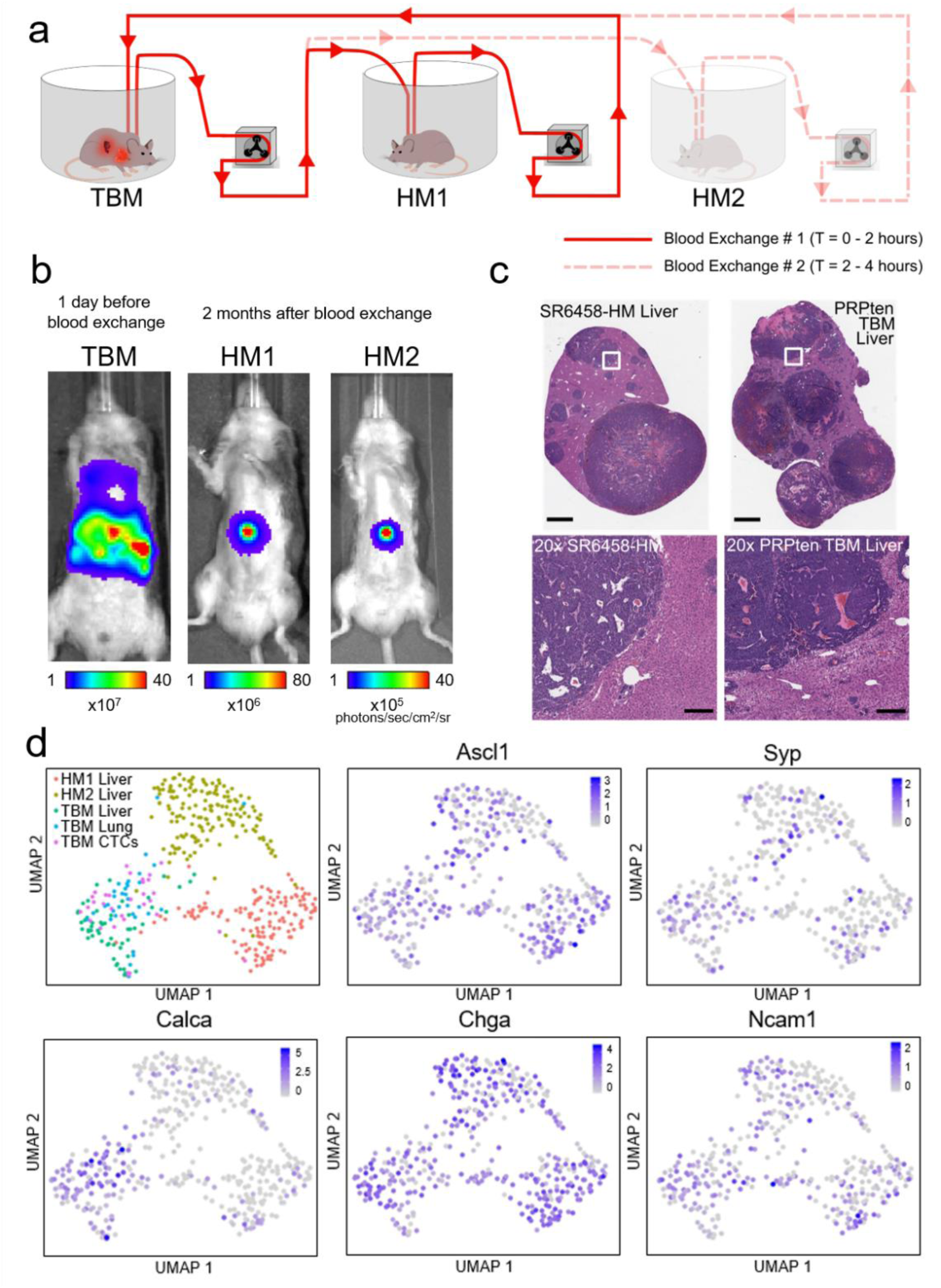
Blood exchange as a method for direct CTC injection for metastasis studies. (**a**) Schematic demonstrating the serial blood exchange experimental setup in which a single TBM was connected to two healthy counterparts, each for two hours. (**b**) Bioluminescent *In Vivo* Imaging System (IVIS) images demonstrating the tumor burden of the donor (TBM) before the blood exchange and the recipient mice (HMs) when liver metastases were detected 2 months after blood exchange. (**c**) Representative H&E-stained liver sections from a blood-exchange recipient mouse (SR6458-HM) and the liver from a tumor bearing mouse (scale bars = 2 mm). Bottom row represent 20x zoomed-in images of the white outlined regions in the top row images (scale bars = 200 µm). (**d**) Uniform Manifold Approximation and Projection (UMAP) plot showing clustering of single-cell transcriptomes of cells across five separate tumor compartments (HM1 liver, HM2 liver, TBM lung, TBM liver and TBM CTCs with 142, 157, 55, 34, and 32 cells per condition respectively) and feature plots showing the expression of 5 known SCLC markers (Ascl1, Syp, Calca, Chga, and Ncam1^41^).

For the late-stage TBMs in our study (5-6 months post tumor initiation) with well***-***established primary lung and metastatic liver burden (**Supplementary Fig. 4b**), the CTC generation rate was 4,000-27,000 CTCs/hour. By subtracting the average weight of healthy (uninfected) PRPten mouse lungs from the average weight of late stage resected lung tumors (**Supplementary Fig. 4c**), we can calculate the approximate weight of the tumor compartment. On the assumption that 1g of SCLC primary tumor contains ∼10^9^ tumor cells^29^ and serves as the main source of these shed CTCs, the ∼16,000 CTCs/hour average generation rate suggests that late-stage SCLCs shed approximately 0.07% of their total tumor cell population per day (or ∼700,000 cells per 1g of tumor tissue). These CTCs exhibited a half-life time in the range of 50-100 seconds. These results demonstrate the utility of the blood-exchange technique for measuring CTC kinetics in a mouse model of SCLC.

### Model Cell Line Circulatory Kinetics differ from CTCs

In order to compare our observed CTC kinetics to traditional studies^2,30^, bolus injections of tumor cell lines were carried out. We collected 25,000 tdTomato-expressing cells prepared from an established murine SCLC cell line that had been generated from murine SCLC tumors isolated from Trp53^fl/fl^; Rb1^fl/fl^; Pten^fl/fl^; Rosa26^LSL-tdTomato/LSL-Luciferase^ mice and propagated *in vitro*^31^. These cells were suspended in 40 µL of either saline or blood and were injected intravenously via the inserted cannula followed by 3-hour scans of the blood with the CTC counter (**Fig. 3a**). In contrast to the CTCs whose behavior is described above, we observed that nearly 98% of the cells cleared from the circulation within seconds (**Fig. 3b-c**). Furthermore, when we repeated the bolus injection experiments using tumor cells that were freshly harvested and dissociated from primary and metastatic tumors from late-stage autochthonous SCLC-bearing mice, we observed a similar rapid clearance of these cells from the circulation, suggesting that this phenomenon was not an artifact of using *in vitro* cultured tumor cell lines. Nonetheless, across all four cell injection experiments, we observed that a small percentage (1-2%) of the injected cells remained detectable in the blood for several hours (**Fig. 3b-c**). These observations provided an initial indication that direct bolus injection of tumor cells into the venous circulation is likely to misrepresent the true dynamics of CTC persistence in the circulation

To examine whether the observed kinetics of clearance following intravenous injection were influenced by the rate at which tumor cells were introduced into circulation, we mimicked the continuous, blood-exchange-based injection rate by using a slow cell-line injection technique (**Fig. 3d**). In these experiments, a small vial containing a freshly prepared suspension of cells, derived from the same SCLC cell line used above, was connected to the venous return line using a T-shaped adaptor to slowly infuse a pre-determined number of cells using a second peristaltic pump. Similar to bolus-cell-line injection experiments described above, slow-injection experiments resulted once again in rapid clearance of cells, represented by the sporadic trajectories of the detected cells (**Fig. 3e** and **Supplementary Fig. 5a**). These trajectories indicate that although the tumor cells were introduced continuously over the duration of each experiment, the majority of the cells cleared extremely rapidly from the circulation. Only a small fraction of injected cells was detected over time with much longer time intervals between subsequently detected cells compared to CTC detection rates. In fact, 7-fold fewer cells were detected with the cell line injection compared to endogenous CTCs during the estimated steady-state intervals (p<0.05, **Fig. 3f** and **Supplementary Fig. 5b-c**). These results further validate the previous bolus intravenous injection results and demonstrate, once again, the stark differences in the circulatory kinetics between endogenous CTCs and intravenously introduced, cultured cells derived from the same tumors.

We next wanted to assess whether the differing half-life times of CTCs spawned by autochthonous tumors and corresponding SCLC cells introduced via the venous circulation was due to differences in cell size, since passage time through a small capillary is strongly size dependent^32–34^. Indeed, our group previously demonstrated that measurements of the buoyant mass of cells in suspension can be used as a proxy for their passage time through a microfluidic constriction^35,36^. In order to compare the buoyant mass of CTCs and their model SCLC cell line, we set out to use the sorting functionality^37^ of the CTC counter to sort and enrich a CTC population from a late-stage SCLC-bearing mouse for subsequent buoyant mass measurements using the suspended microchannel resonator (SMR) platform^38,39^. As a control, a viable SCLC cell line population was spiked into a blood sample collected from a healthy mouse (1,000 tdTomato-expressing cells per 100 µL of blood), after which single cells were sorted and enriched using our CTC counter for mass measurements (**Supplementary Fig. 6a**). Interestingly, both cell populations had a similar buoyant mass distribution (p = 0.3, Mann-Whitney-Wilcoxon non-parametric test, **Supplementary Fig. 6b**), suggesting that differing sizes of the cells in these two populations could not be invoked to explain their different dwell times in the circulation. This would suggest, as an alternative, that the intrinsic biological properties of these two populations of cells could be the dominant determinant**s** of the unique circulatory kinetics for each population.

We proceeded to examine whether these two cell populations did indeed exhibit distinct biological properties contributing to the differing lifetimes in the circulation. To do so, we employed single-cell RNA-Sequencing (scRNA-Seq) to profile the SCLC cell line along with a group of CTCs sorted from a terminal blood sample of an autochthonous SCLC-bearing mouse. Principal Component Analysis (PCA) on highly variable genes revealed stark differences in the two populations, with CTCs associated with a positive PC1 and negative PC2, and cell line associated with a negative PC1 and positive PC2 (**Supplementary Fig. 6c**). In order to explore the biological drivers of this separation, we performed correlation analysis between the principal components and expression of Gene Ontology genesets. This analysis revealed that PC1 was positively correlated with genesets for epithelial-to-mesenchymal transition (EMT) and cytoskeletal organization, and was negatively correlated with translation and cell cycle genesets (**Supplementary Fig. 6d**). This indicates that EMT, and cytoskeleton markers are associated with CTCs as opposed to the *in vitro* cell line counterpart, which had higher expression of cell cycle-related genes. These findings also held for PC2, which correlated positively with translation genesets and negatively with cytoskeletal organization and cell stress genesets. Further experiments will be needed to validate these findings and explore how these transcriptomic variations influence the behavior of the corresponding cells in the circulation as well as the locations and phenotypes of the distant tumors they form.

### Blood Exchange as a Method for Generating Metastases in Naïve Healthy Mice

In two separate experiments, we observed that the direct introduction of as few as 4,000-7,000 CTCs via blood exchange over a few hours (representing only 1-2% of the average daily shed CTCs) was capable of generating liver and intestinal metastases in healthy recipient mice within two to three months (**Supplementary Fig. 7a**). Interestingly, the newly developed liver metastatic lesions were also capable of shedding CTCs as confirmed by the detection of tdTomato-expressing CTCs in the terminal blood of those mice (**Supplementary Fig. 7b**).

We next set out to further investigate the ability of our method to study the metastatic propensity of CTCs introduced into a healthy mouse by blood exchange. Two healthy immunocompetent mice (IDs: HM1 and HM2) were sequentially connected to a late-stage SCLC GEMM, and approximately 8,000 CTCs were introduced into each healthy mouse over 2 hours of blood exchange (**Fig. 4a**). After the mice were disconnected, the TBM was sacrificed in order to collect individual tumor cells from the lung and liver as well as CTCs from the blood. The healthy recipient mice were longitudinally monitored over several weeks using *in vivo* Bioluminescence Imaging (BLI) for tumor development. Both recipient mice developed liver macrometastases approximately 2 months post blood exchange (**Fig. 4b**). Interestingly, macrometastases were observed in the liver rather than the lung, even though the CTCs introduced originated from a lung tumor and, having entered via entry through the right jugular vein, encountered the microvessels of lung in the healthy mouse as the first major impediment to remaining in the circulation. This observation is consistent with the dissemination pattern of the SCLC GEMM as well as clinical findings in which liver metastases are reported as a common dissemination site in SCLC patients^40^.

To further validate that the developed tumors in the HMs resemble those of the donor mouse, HM1 and HM2 were sacrificed following tumor detection for histological and transcriptomic analysis. Findings of hematoxylin and eosin (H&E)-stained liver sections from one of the HM lesions confirmed the histological features of a SCLC metastatic liver tumor (**Fig. 4c**). We then performed scRNA-Seq analysis on cells prepared from the tumor samples and terminal CTCs harvested from the one TBM donor, as well as the tumors collected from the two HM recipients. These cells exhibited high quality control, indicated by transcript and gene counts, as well as high purity, indicated by the expected high expression of EpCAM and tdTomato and low expression of blood cell genes including protein tyrosine phosphatase receptor type C (CD45) and platelet factor 4 (Pf4) (**Supplementary Fig. 8**, see scRNA-Seq in *Methods*). Differential expression comparisons between the donor lung tumor and all three liver tumors (TBM, HM1, and HM2) revealed a large number of overlapping genes (**Supplementary Fig. 9a**). 20% of the differentially expressed genes were consistent in at least 2 of the liver tumors, and 5% were consistent in all 3. Although mouse-specific clustering was observed in the Uniform Manifold Approximation and Projection (UMAP) plot (**Fig. 4d, Supplementary Fig. 9b**), this may be due to local microenvironment effects, different stages of tumor development, or the presence of multiple independently evolving tumors within each mouse^27^. Furthermore, a set of 5 canonical SCLC genes were used to verify the tumor transcriptionally as SCLC^41^ (**Fig. 4d**). Comparable expression of this geneset across tumor compartments further validates these metastatic lesions as SCLC metastases. Taken together, these findings confirm that the blood exchange technique can be used to create metastatic lesions in healthy recipient mice. Future experiments will be needed to robustly validate these findings, and thoroughly explore the differences between donor and recipient tumors.

## Discussion

Here we demonstrate a novel, pre-clinical blood-exchange method to study the circulatory kinetics of CTCs in mice. Examination of the shedding rate was last demonstrated using an artificial rat model nearly 45 years ago, when mammary adenocarcinoma CTCs were reported to be shed from solid tumors at a daily rate of 3.2 to 4.1×10^6^ cells per gram of tissue^43^, which is ∼5 times higher than our own estimates (∼700,000 per gram of tissue). Previous estimates of CTC half-life in the mouse circulation have varied from seconds^2,44^ to minutes^14^ to hours^45^ using a variety of detection techniques. In clinical settings, these have relied on CTC detection rates in the hours following tumor resection surgery^45^, while pre-clinical animal models primarily have employed bolus injections of fluorescent tumor cell lines tracked by in-vivo flow cytometry (IVFC)^46,47^. Although longer half-life times of 30-60 minutes have been previously reported, the measurements were based on only a small fraction of the injected cell population, as the majority of these bulk-injected cells were cleared almost instantaneously^2,3^. We contrast this earlier work with the half-life time estimates obtained from our blood-exchange technique, which involved naturally shed, endogenous CTCs and relied on the positive detection of their real-time entry and exit rates from the circulatory systems of the two connected mice. In contrast to CTCs, when we mimicked these experiments using cell lines in a slow-injection configuration, we observed kinetics that were consistent with previously reported cell line injection experiments. Our results raise the question of whether the previous estimates of CTC dwell time in the circulation were strongly influenced in an artifactual manner by the experimental procedures employed.

The blood-exchange technique discussed here also introduces a new method for generating metastases using true CTCs. Traditional methods either use cell lines or require the isolation, *ex vivo* expansion, and characterization of specific subpopulations of CTCs endowed with metastatic potential prior to their inoculation in animal models^14,20–22,48^. In our method, CTCs from mice bearing primary and metastatic lesions remain in their original blood and directly enter the circulatory system of the healthy immunocompetent recipient mouse without the need of enrichment or intervening culturing steps that may affect their viability and profoundly change their biological traits. Importantly, the presently employed method may extend the time window for studying metastatic disease, particularly in tumor models in which host mice succumb to primary burden before fully-developed metastases can be analyzed^40^. Further studies will be needed to validate the utility of this technique in other cancer types.

In summary, the reported blood-exchange technique and resulting CTC kinetics data can lead to accurate identification of the rate-limiting steps in the blood transport phases of the metastatic cascade. We also envision that our blood-exchange technique can be used to directly and controllably exchange other blood components and study trafficking dynamics of immune cells in various biological contexts within immunology, cancer biology, and aging. Because our blood-exchange technique can be used continuously and longitudinally, it can potentially reveal temporal kinetics that occur on the order of minutes, hours, or days and hence may assist in establishing suitable time windows for maximizing therapeutic efficacy.

## Methods

### The Blood-Exchange Optofluidic Platform

The platform is a modified Mouse CTC Sorter^37^ that consists of three major subsystems: a microfluidic device, an optical detection system, and a computer control system. The microfluidic chip is designed with one inlet to a 1 cm-long microfluidic channel (300 × 45 µm) that bifurcates into two channel outlets (90° apart); one for returning the blood to the mouse and the other for collecting the sorted CTC-containing blood sample. The microfluidic device comprises two polymer layers bonded to a glass base layer. The bottom polymer layer implements the pneumatically activated valve structures that control the fluid path through channels in the top layer.

In normal operation, the collection valve is closed and the return valve is open and CTCs are just enumerated as they pass through the system. If a CTC is to be collected, the return valve is closed, and the collection valve opens momentarily to deflect a small amount of blood containing the CTC. At a flow rate of 30 µL/min, the average sort volume is 127±47 nanoliters/CTC.

### Device Fabrication

The fabrication of the microfluidic chip for the CTC counter, as reported previously^37^, starts with standard soft lithographic techniques on two four-inch wafers. A single layer of photoresist (SU8 2050, Microchem, Newton, MA) is patterned to create the pneumatic channels on the valve actuation wafer. For the blood flow channel, AZ9260 positive resist was exposed, developed, and then reflowed at 120° C for 10 minutes to create the rounded channel profile necessary for a complete valve seal. Once the master mold is fabricated for both the actuation and flow channel layers, a mixture of PDMS (Polydimethylsiloxane) and its curing agent (SYLGARD 184 A/B, Dowcorning, Midland, MI, USA) at a 10:1 ratio was spun on top of the actuation wafer to a thickness of 50 µm and baked in an oven set to 65° C for 3 hours. For the flow channel layer, the mixture was poured to a thickness of ∼1 cm and cured at 65° C for 3 hours. Afterwards, the flow channel layer was peeled off and punched with a 0.75 mm puncher (Harris Uni-Core, Ted Pella Inc., Reading, CA) to define the inlet and outlets to and from the flow channel, respectively, and diced to prepare for bonding. The flow channel devices and the actuation layers were then treated with oxygen plasma (100 watt, 1 ccm, 140 torr, 10 seconds). Next, the flow layer was aligned to the actuation layer and baked in an oven at 60°C. After 15 minutes, the assembled PDMS layers were peeled off the wafer and punched with a 0.75 mm puncher to define inlets to the actuation channels. The assembled PDMS layers were treated with oxygen plasma (100 watt, 1 ccm, 140 torr, 10 seconds) for irreversible bonding to a glass slide (Fisherbrand 1×3”, Fisher Scientific, Pittsburgh, PA). Prior to flow experiments, the device was aligned to project the two laser lines across the flow channel 8 mm away from the valve actuation region. The device was then primed with Heparinized-Saline (diluted to 100 USP units per mL, NDC 25021-400-30) to prevent clotting within the microfluidic channel.

### CTC Counter Optical Setup

The CTC counter consists of two optical trains aligned vertically. The top optical train also consists of a dichroic filter to reflect the excitation beam (532 nm) onto the sample and transmit the emitted signal (greater than 532 nm). The second dichroic mirror and longpass filters, placed constantly directly above the detection region during normal operation, pass a filtered fluorescence signal to the PMT (Hamamatsu H10722-20) by further blocking the 532 laser line with a notch filter. A 90:10 (T:R) beam splitter is added before the PMT to allow for imaging of the illumination region for device alignment purposes during initial experimental set up. The bottom optical train, for verifying proper microvalve functionality and sorting mechanism of the chip, consists of similar components to the top optical train but shifted laterally by approximately 1 cm.

### Mice

All animal-based procedures were approved by the Massachusetts Institute of Technology Committee on Animal Care (CAC), Division of Comparative Medicine (DCM). All mice were maintained on a mixed C57BL/ 6;129/Sv background. The Trp53^fl/fl^; Rb1^fl/fl^; Pten^fl/fl^; Rosa26^LSL-Luciferase/LSL-Luciferase^(PRP-L/L) mouse model of SCLC has been described previously^27^. Rosa26^LSL-tdTomato/LSL-tdTomato^ mice were obtained from Jackson Laboratories (Gt(ROSA)^26Sortm14(CAG-tdTomato)Hze^) and crossed into the PRP-L/L model to obtain Trp53^fl/fl^; Rb1^fl/fl^; Pten^fl/fl^; Rosa26^LSL-tdTomato/LSL-Luciferase^ mice (genotype of all TBM and HM used in this study). Tumors in TBMs are initiated by the delivery through the trachea of 2×10^8^ plaque forming units (p.f.u.) of adenovirus expressing Cre recombinase under the control of a CGRP promoter (Ad5-CGRP-Cre), as previously described^49^. Adenoviral stocks were purchased from the Viral Vector Core Facility at the University of Iowa Carver College of Medicine.

### Cannulation Surgery

All animal-based procedures were approved by the Massachusetts Institute of Technology Committee on Animal Care (CAC), Division of Comparative Medicine (DCM). Candidate mice for the arteriovenous shunt surgery were identified by in vivo bioluminescence imaging using the IVIS Spectrum In Vivo Imaging System (PerkinElmer). As reported previously^37^, catheters are inserted into the right jugular vein and the left carotid artery and are externalized using standard cannulation surgical techniques in anesthetized mice.

### Cell Line Injection Studies

A murine SCLC cell line (AF3291LN) was generated from SCLC tumors isolated from Trp53^fl/fl^; Rb1^fl/fl^; Pten^fl/fl^; Rosa26^LSL-tdTomato/LSL-Luciferase^ mice as previously described^31^. For bolus cell line injections, a population of at least 1 million cells is harvested from a flask, rinsed with saline, and centrifuged at 1000 rpm for 5 minutes. Pelleted cells were then re-suspended in saline and counted using the CTC counter to dilute the sample to a dosing concentration of 25,000 cells per 40 µL.

For slow injection experiments, similar washing steps were executed but the final cell density is reduced to a desired concentration between 3,000 and 20,000 cells/mL. The cell-suspension vial is connected to a second peristaltic pump that infuses the cells intravenously through a T-adaptor at a fixed flow rate of ∼2 µL/min. Throughout the injection experiment, the cell suspension vial is replaced with a new, well-mixed cell-suspension vial every 15 minutes to ensure a consistent infusion of viable cells. To determine the approximate injection count over time (blue trajectories in **Fig. 3e** and **Supplementary Fig. 5a**), a small sample from each freshly-prepared vial is taken and counted on a separate CTC counter.

### SMR Experiment

The silicon-based SMR chip is the core component of the SMR system and consists of a sealed microfluidic channel that runs through the interior of a cantilever resonator^50^. As a cell in suspension flows through the cantilever, it transiently changes the cantilever’s resonant frequency in proportion to its mass. A fluorescent readout was integrated with the SMR to detect the tdTomato-expressing CTCs in a heterogenous sample. In this modified SMR system, an excitation laser beam is focused into a 500 µm line and aligned across the inlet bypass channel of the SMR chip (**Supplementary Fig. 6a**). Emitted signal from a tdTomato-expressing CTC passing under the laser line is detected by a PMT (similar to the real-time CTC counter) to induce an automated fluidic direction change that slowly loads the cell into the cantilever section for mass measurements. Prior to their mass measurement, both samples (CTCs and enriched cell line) are re-suspended in growth media (DMEM with 10% FBS).

### Dissociation of Tumor Samples for Single-Cell RNA-Sequencing Analysis

Primary lung and metastatic liver tumors from tumor-bearing animals were dissected, dissociated into single cells using dissociation kits according to the manufacturer’s protocol (preserving all surface epitopes, Miltenyi Biotec #130-095-927), then stained with APC-conjugated antibodies against CD45 (eBioscience #17-0451-83). tdTomato-expressing, APC-negative cells were single-cell sorted by FACS into TCL buffer (QIAGEN) containing 1% 2-mercaptoethanol, and then frozen at -80°C for downstream processing for scRNA-Seq. According to the manufacturer, the dissociation protocols

### CTC Enrichment for Single-Cell Analyses

CTC-enriched blood sorted from the CTC counter is further purified through sequential dilution. The pooled CTCs (in approximately 100 nL/CTC) are diluted to 500 µL in cell culture media (DMEM + 10% FBS). The diluted blood is run through the CTC counter again, and fluorescent CTCs are re-sorted in approximately 100 nL of media and re-diluted. After 3 dilution steps in media, the CTCs are diluted a final time to 500 µL in RNase-free PBS, run through the CTC counter, and collected into a PCR tube containing 7 µL of 2×TCL lysis buffer (Qiagen) with 2% v/v 2-mercaptoethanol (Sigma). The samples are immediately frozen on dry ice and subsequently stored at −80° C until library preparation and sequencing. The complete process results in a typical dilution between 3.9×10^5^ and 6.3×10^6^, depending on the number of detected CTCs.

### Single-Cell RNA-Sequencing Sample Preparation

CTC and primary tumor samples in TCL supplemented with 1% (v/v) 2-mercaptoethanol buffer were processed through Smart-Seq2 as previously described^37,51^. Briefly, cellular nucleic acids from each lysed single cell were extracted from TCL lysis buffer using a 2.2x (v/v) RNA SPRI (RNA-clean AMPure beads, Beckman-Coulter). After, we performed reverse transcription was Maxima enzyme (Thermo scientific), and then PCR using KAPA Hotstart Readymix 2x kit (KAPA biosystems). Following quantification and quality control analysis by Qubit DNA quantification (Thermoscientific) and tape station (Agilent), the post-PCR whole transcriptome amplification (WTA) products from each single cell were transformed into sequencing libraries using a Nextera XT kit (Illumina) and unique 8-bp DNA barcodes. cDNA libraries were pooled, quantified, and sequenced on an Illumina NextSeq 500 to an average depth of 1.2M reads/cell.

### Analysis of Raw Sequencing Data

Following sequencing, BCL files were converted to merged, demultiplexed FASTQs. Paired-end reads were mapped to mm10 mouse transcriptome (UCSC) with Bowtie 2. Expression levels of genes were log-transformed transcript-per-million (TPM[i,j]) for gene i in sample j, estimated by RSEM in paired-end mode. For each cell, we enumerated genes for which at least one read was mapped, and the average expression level of a curated list of housekeeping genes. We excluded from analysis profiles with fewer than 500 detected genes or below 375,000 total reads (**Supplementary Fig. 8a**), though downstream results were consistent with more and less stringent cutoffs. Principal component analysis (PCA) was performed on all variable genes except Gm, RP, and Hb genes as initial results indicated a dominant method of sorting signature within the dataset driven by these genes (CTC counter for sorting CTCs and fluorescence-activated cell sorting (FACS) for sorting cell line, lung, and liver samples).

### PC Correlation Analysis

To understand correlative effects of metastasis-related genesets on the PC separation of CTCs vs cell line (**Supplementary Fig. 6c**), module scores were added for each cell for several Gene Ontology genesets [Epithelial to Mesenchymal Transition (GO:0001837), Cytoskeleton Organization (GO:0007010), Regulation of Cell Stress (GO:0080135), Fluid Shear Stress (GO:0034405), Cell Cycle (GO:0007049), Cytoplasmic Translation (GO:0002181), Translation Initiation (GO:0006413)] using the AddModuleScore function in Seurat. Then, Pearson Correlation coefficients were calculated between the module scores for each geneset and both PCs 1&2. Finally, the resulting R values were visualized using the Complexheatmap package (**Supplementary Fig. 6d**), and p values were calculated using R value and number of samples. To confirm that PC separation was not driven by quality variations, cutoffs were modulated to ensure comparable levels of quality control metrics. The same trends in PC correlation to GO genesets was found, and quality control metrics had no significant correlation.

### UMAP Analysis

Variable genes across the tumor compartments (for blood exchange experiment) or cell samples were calculated and principal component analysis was performed using Seurat. The first 10 principal components were used in Seurat (determined by the Elbow Plot) to run UMAP analysis, though the clustering was robust for varying numbers of dimensions used. Further results were visualized using the FeaturePlot and DoHeatmap functions in Seurat.

### Differential Expression Analysis

Since the 3 liver tumors clustered independently, we used differential expression analysis to explore the varying genes. For each liver sample, we performed differential expression relative to the TBM primary lung sample with Seurat’s built-in single-cell differential expression tool using a bimodal distribution model. Genes with average logfold-change > 0.6 and Bonferroni adjusted p value < 0.05 were selected, though similar trends were seen for both more and less stringent cutoffs, and overlapping genes were visualized using a Venn Diagram.

## Acknowledgments

We thank P. Winter, T. Meittinen, G. Katsikis, L. Mu, L. Li, B. Eskiocak, A. Cruz, D. Benjamin, R. Hynes, and C. Jin for helpful discussions regarding different aspects of the blood exchange experimental set up. We thank the Koch Institute Swanson Biotechnology Center for technical support, specifically the Animal Imaging & Preclinical Testing Core Facility, the Flow Cytometry Core Facility, and the Hope Babette Tang (1983) Histology Facility. This work was supported, in part, by the Thomas and Sarah Kailath Fellowship (to B.H.); the Ludwig Center for Molecular Oncology Graduate Fellowship (to B.H.); the A*STAR (Agency for Science, Technology and Research, Singapore), and National Science Scholarship (to S.R.N.). We acknowledge funding from the Ludwig Center for Molecular Oncology (S.R.M.), the Cancer Systems Biology Consortium U54 CA217377 and the Koch Institute Support Grant P30 CA14051 from the NCI (S.R.M.). A.K.S. was supported by the Pew-Stewart Scholars Program for Cancer Research, a Sloan Fellowship in Chemistry, and the NIH (1DP2GM119419, 1U54CA217377 and 2RM1HG006193).

