## Supplementary Figures for "Measuring kinetics and metastatic propensity of CTCs by blood exchange between mice"

[1]Department of Electrical Engineering and Computer Science, Massachusetts Institute of Technology, Cambridge, Massachusetts 02139, USA
[2]David H. Koch Institute for Integrative Cancer Research, Massachusetts Institute of Technology, Cambridge, MA 02142, USA
[3]Harvard-MIT Department of Health Sciences and Technology, Institute for Medical Engineering and Science, Massachusetts Institute of Technology, Boston, MA 02142, USA
[4]Department of Oncology, Hematology and Bone Marrow Transplantation with Section Pneumology, Hubertus Wald Comprehensive Cancer Center Hamburg, University Medical Center Hamburg-Eppendorf, Martinistrasse 52, 20246, Hamburg, Germany.
[5]Department of Tumor Biology, Center of Experimental Medicine, University Medical Center Hamburg-Eppendorf, Hamburg, Germany.
[6]Department of Mechanical Engineering, Massachusetts Institute of Technology, Cambridge, MA 02139
[7]Department of Biology, Massachusetts Institute of Technology, Cambridge, MA 02139
[8]Department of Biomedical Engineering, Wentworth Institute of Technology, Boston, MA 02115
[9]Whitehead Institute for Biomedical Research, Cambridge MA 02142
[10]Department of Chemistry, Massachusetts Institute of Technology, Cambridge, MA 02139;
[11]Institute for Medical Engineering and Science, Massachusetts Institute of Technology, Cambridge, MA 02139;
[12]Broad Institute of MIT and Harvard, Cambridge, MA 02142;
[13]Ragon Institute of MGH, MIT and Harvard University, Cambridge, MA 02139;
[14]Department of Immunology, Massachusetts General Hospital, Boston, MA 02114
[15]Howard Hughes Medical Institute, Massachusetts Institute of Technology, Cambridge, MA 02139
[16]Department of Biological Engineering, Massachusetts Institute of Technology, Cambridge, MA 02139;
*co-first author
†corresponding author

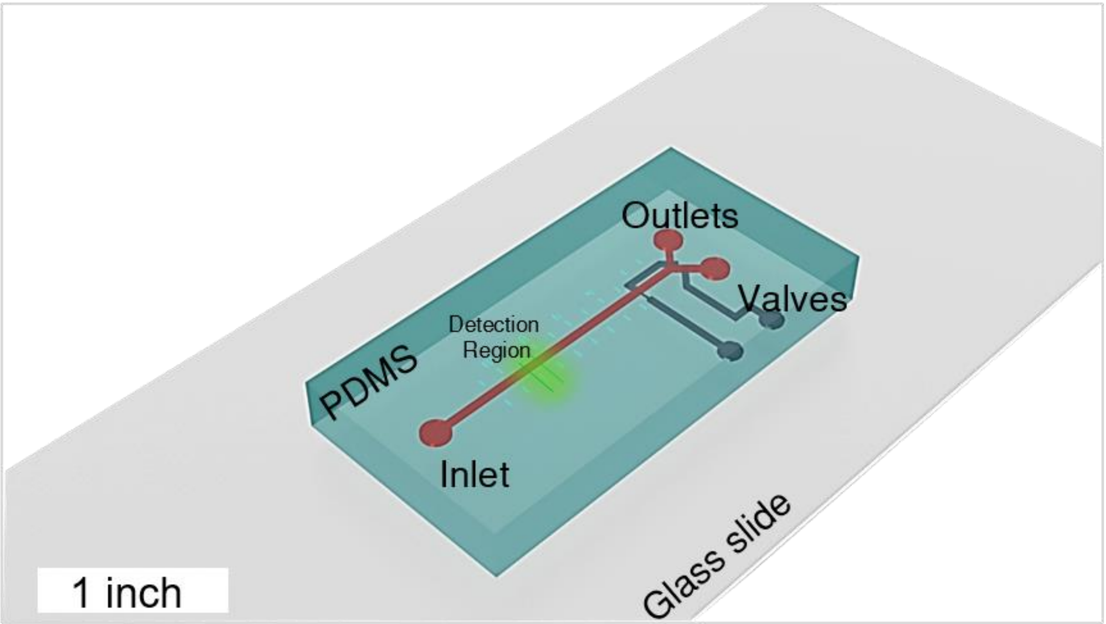


**Supplementary Fig. 1 |** Three-dimensional rendering of the PDMS-based CTC counter microfluidic device for counting or sorting CTCs in real-time. CTCs pass through dual excitation laser lines focused near the inlet of the device (Detection Region). Two outlets for either returning the blood to the mouse or collecting the CTC when the sorting functionality is activated. Microvalves near the outlet control blood flow out of the microfluidic chip.

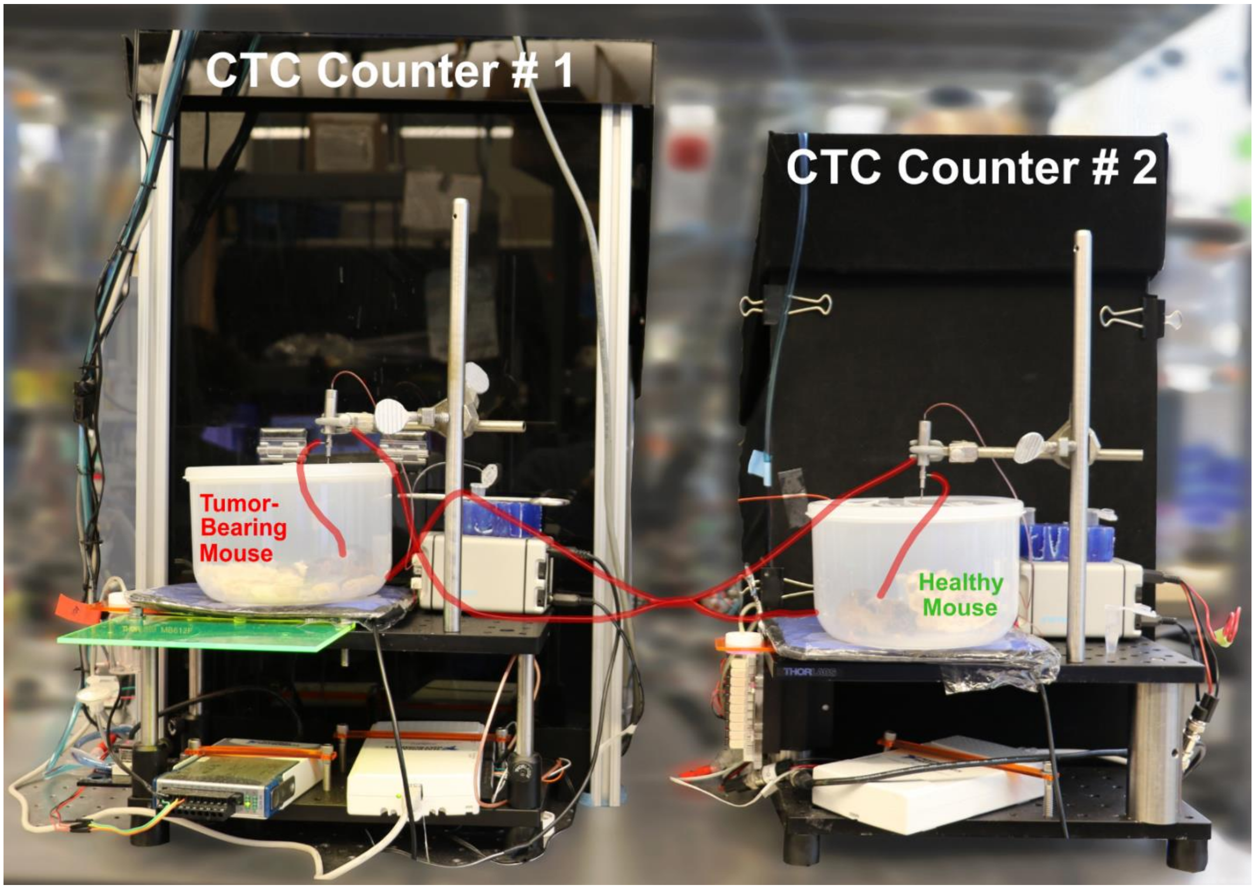


**Supplementary Fig. 2 |** A front view demonstrating the blood-exchange set up. Two mice (TBM and HM) with tethered externalized carotid artery and jugular vein catheters move freely in a small container. Peristaltic pumps withdraw blood from the carotid artery into the inlet of the optofluidic CTC counter (black box behind each mouse container). Return blood catheters (highlighted in red) transport blood back to the jugular vein of the other mouse to complete the blood flow loop without loss. Both systems are controlled by two separate LabVIEW controllers and the entire system is suspended on optical tables to isolate vibrational noise.

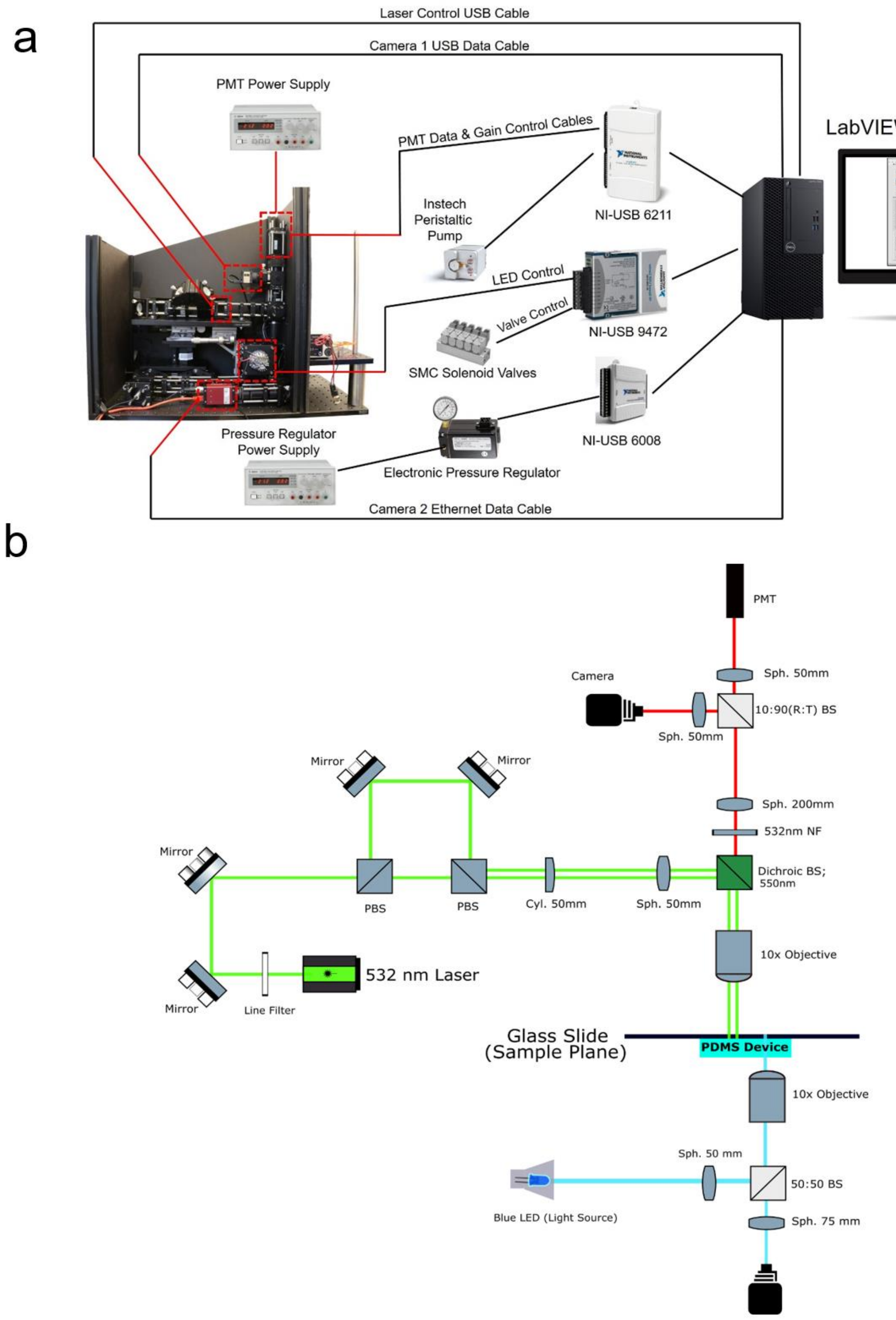


**Supplementary Fig. 3 |** Main hardware and optical components of the CTC counter. (**a**) Side-view image of the CTC counter with visual illustrations of the different interfacing components for data flow. (**b**) Schematic of the optical components of the CTC counter. A single excitation source (532 nm Laser) on the top optical train is filtered, split, and then projected on the sample plane for CTC detection within the PDMS-based microfluidic device. Emitted signal is filtered prior to being focused on a PMT for signal detection. Bottom optical train is used for real time monitoring of blood flow at the outlet region.

a

| TBM ID | Exp. # | Q (mL/min) | V (mL) | $r_1$ (cells/min) | $r_1$ SD | $r_2$ (cells/min) | $r_2$ SD | $r_{gen}$ (CTCs/hour) | $r_{gen}$ Err | $t_{1/2}$ (sec) | $t_{1/2}$ Err |
|---|---|---|---|---|---|---|---|---|---|---|---|
| SR7401 | **1** | 0.06 | 1.57 | 5.90 | 0.82 | 0.49 | 0.03 | 4282.02 | 1232.03 | 97.28 | 15.85 |
| SR7464 | **2** | 0.06 | 1.51 | 17.10 | 0.61 | 1.22 | 0.10 | 14352.69 | 1599.76 | 80.08 | 7.92 |
| SR7402 | **3** | 0.06 | 1.59 | 11.85 | 0.51 | 0.48 | 0.09 | 17624.38 | 3531.58 | 46.10 | 8.88 |
| SR6458 | **4** | 0.06 | 1.52 | 17.67 | 0.20 | 0.97 | 0.13 | 19253.09 | 2638.61 | 61.24 | 8.73 |
| SR7391 | **5** | 0.06 | 1.46 | 25.20 | 0.40 | 1.39 | 0.09 | 27380.58 | 1950.07 | 59.07 | 4.10 |

b

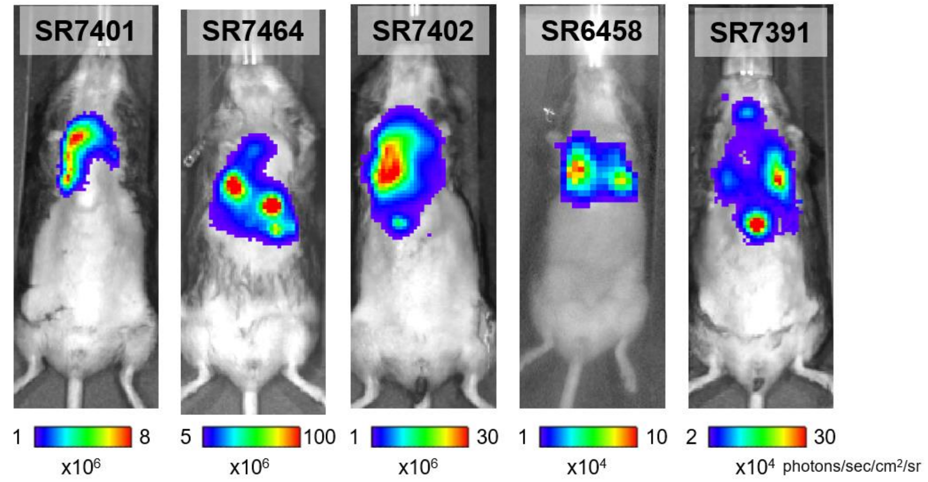


c

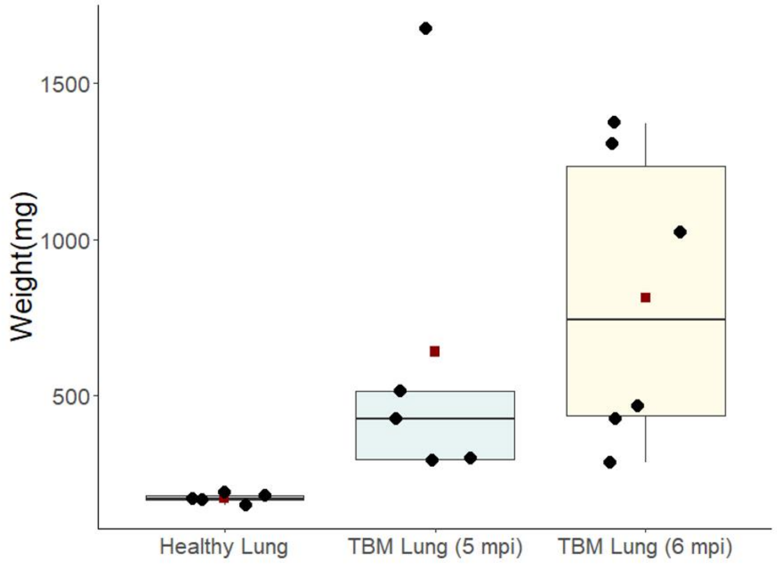


**Supplementary Fig. 4 |** (**a**) Table summarizing the main parameters extracted from the CTC counts during the blood exchange experiments. (**b**) Bioluminescence IVIS images of each of the TBMs demonstrating a visible tumor burden in the days or weeks prior to their blood-exchange experiments with healthy counterparts. (**c**) Boxplot of lung weights collected from five (black) healthy and ten tumor-bearing (green for 5 mpi and yellow for 6 mpi) mice. mpi- months post infection. Box denote the interquartile range with red square means.

a

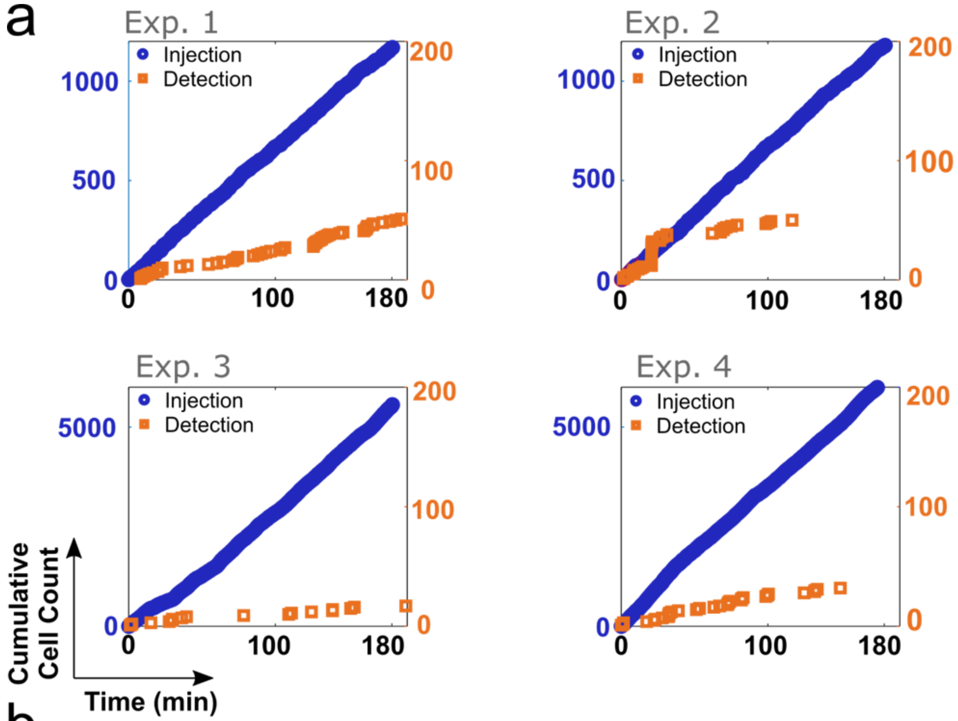


b

| SS Interval (min) | Blood Exchange Detection Fraction (%) | | | | | Slow Injection Detection Fraction (%) | | | |
|---|---|---|---|---|---|---|---|---|---|
| | Exp. 1 | Exp. 2 | Exp. 3 | Exp. 4 | Exp. 5 | Exp. 1 | Exp. 2 | Exp. 3 | Exp. 4 |
| 30 | 9.70 | 7.58 | 4.90 | 5.58 | 6.10 | 4.55 | 0.00 | 0.19 | 0.00 |
| 60 | 8.21 | 7.47 | 4.78 | 6.14 | 5.94 | 5.18 | 0.00 | 0.24 | 0.24 |
| 90 | 7.33 | 6.39 | 3.75 | 7.15 | 6.58 | 4.43 | 0.63 | 0.23 | 0.27 |
| AVG | 8.41 | 7.15 | 4.48 | 6.29 | 6.21 | 4.72 | 0.21 | 0.22 | 0.17 |
| SD | 1.20 | 0.66 | 0.63 | 0.80 | 0.33 | 0.40 | 0.36 | 0.03 | 0.15 |

c

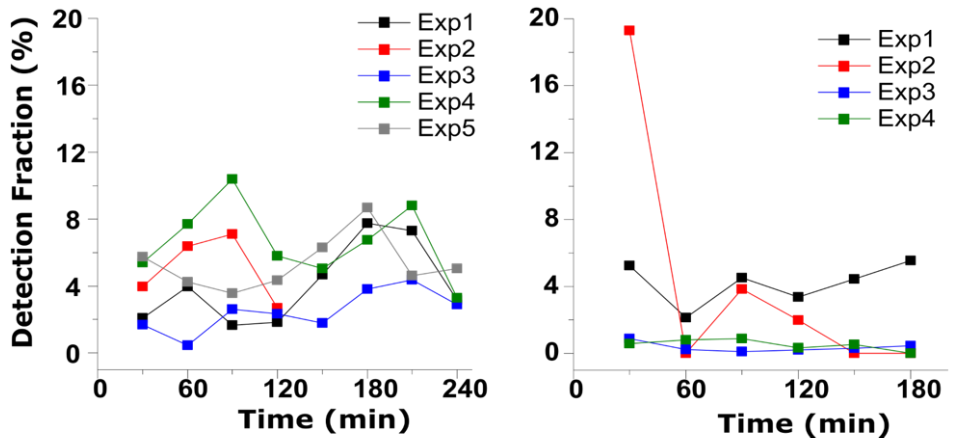


**Supplementary Fig. 5 | (a)** Cumulative injected (blue) and detected (orange) cell counts from four slow-injection experiments. Fluorescent SCLC cell were suspended in saline at pre-determined concentrations and then slowly introduced into the circulatory system of four different healthy recipient mice. Cell counts in blood are monitored for three hours during the slow injection (orange). (**b**) Table summarizing the fraction of detected cells to total injected cells (“Detection Fraction”) in the assumed steady-state intervals at the end of each of the nine experiments (five blood exchange and four slow-injection experiments). (**c**) Detection fraction over 30-minute intervals throughout the blood-exchange (left) and slow-injection (right) experiments.

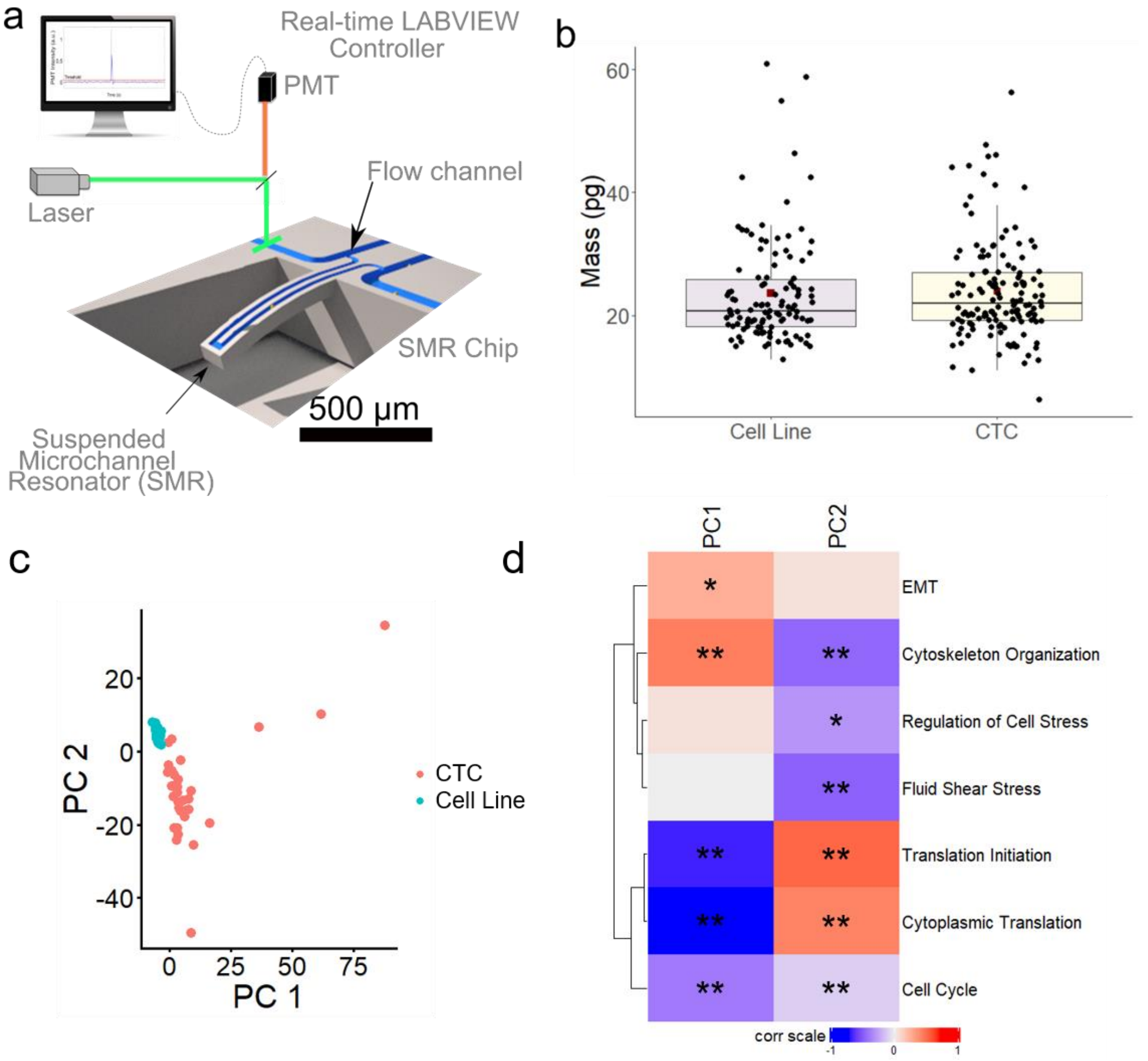


**Supplementary Fig. 6 |** (**a**) Schematic demonstrating the main components of the suspended microchannel resonator (SMR) platform that was designed and built for measuring the buoyant mass of single fluorescent cells in heterogeneous samples. A laser excitation source is aligned across a bypass channel upstream of the mass sensor (cantilever). Upon the detection of a target fluorescent cell or CTC, an automated active-loading system is triggered to allow for the flow to be directed through the cantilever for their mass measurement. (**b**) Box plot demonstrating the mass distribution of detected CTCs and their model SCLC cell line. No significant difference was observed between the two groups ($p = 0.3$, Mann-Whitney-Wilcoxon non-parametric test). (**c**) Principal Component Analysis (PCA) plot showing separation of scRNA-Seq data between true CTCs and the cell line population. (**d**) Heatmap showing correlation coefficients between the PCs 1&2 with expression of select Gene Ontology genesets. Color represents Pearson Coefficient, R. *$p<.01$; **$p<.0001$.

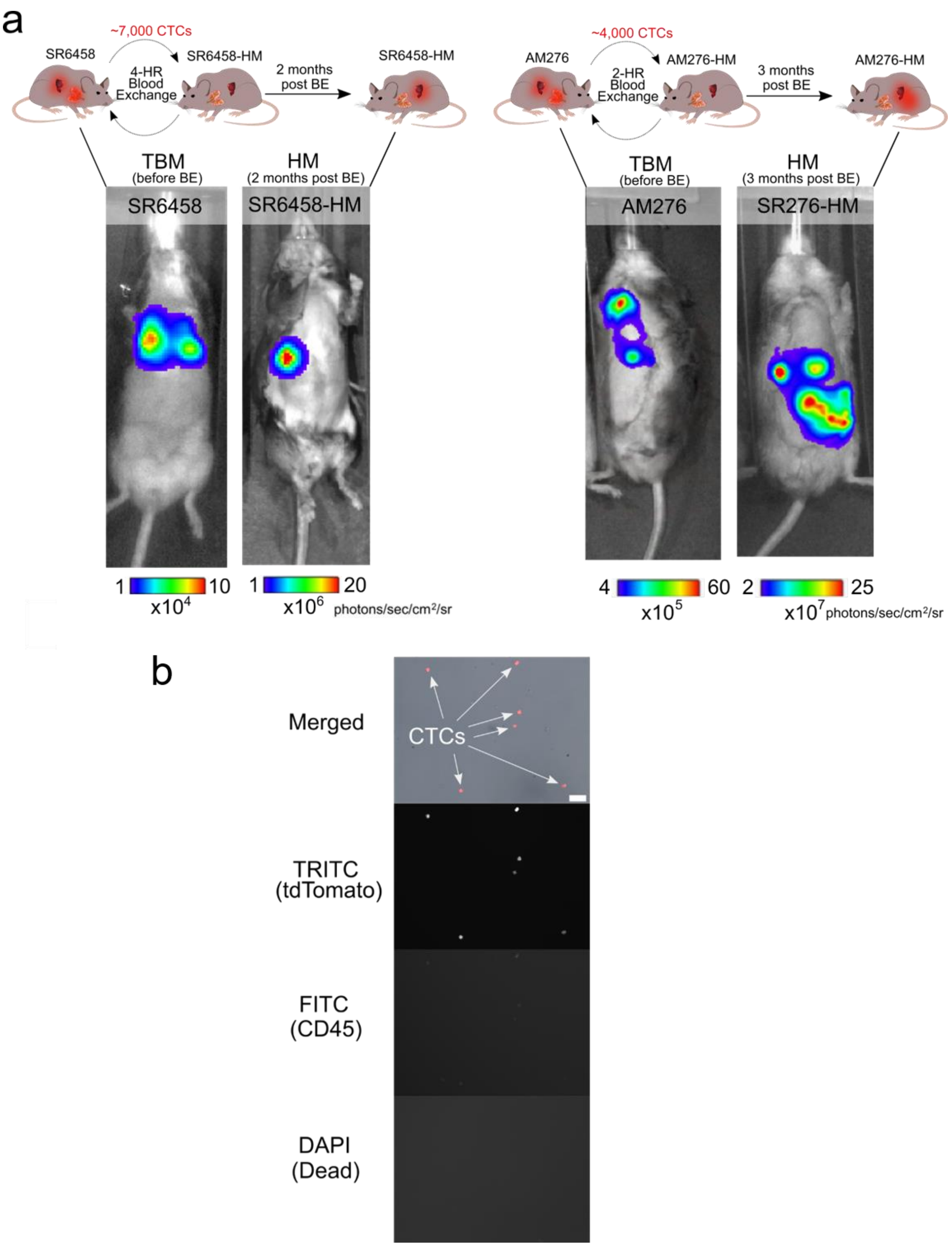


**Supplementary Fig. 7 |** (**a**) Bioluminescence IVIS images of two TBMs prior to blood exchange experiments and two originally healthy counterparts two months after the blood-exchange experiments. Superimposed rainbow-colored regions represent the detectable lung and liver burden in the TBMs and the liver/intestinal burden in the HMs. (**b**) 10x microscopy images of different fluorescent channels of single-cell sorted CTCs into a well of a glass-bottom multi-well plate from the terminal blood of the HM (two months after the blood exchange experiment, scale bar = 20 μm).

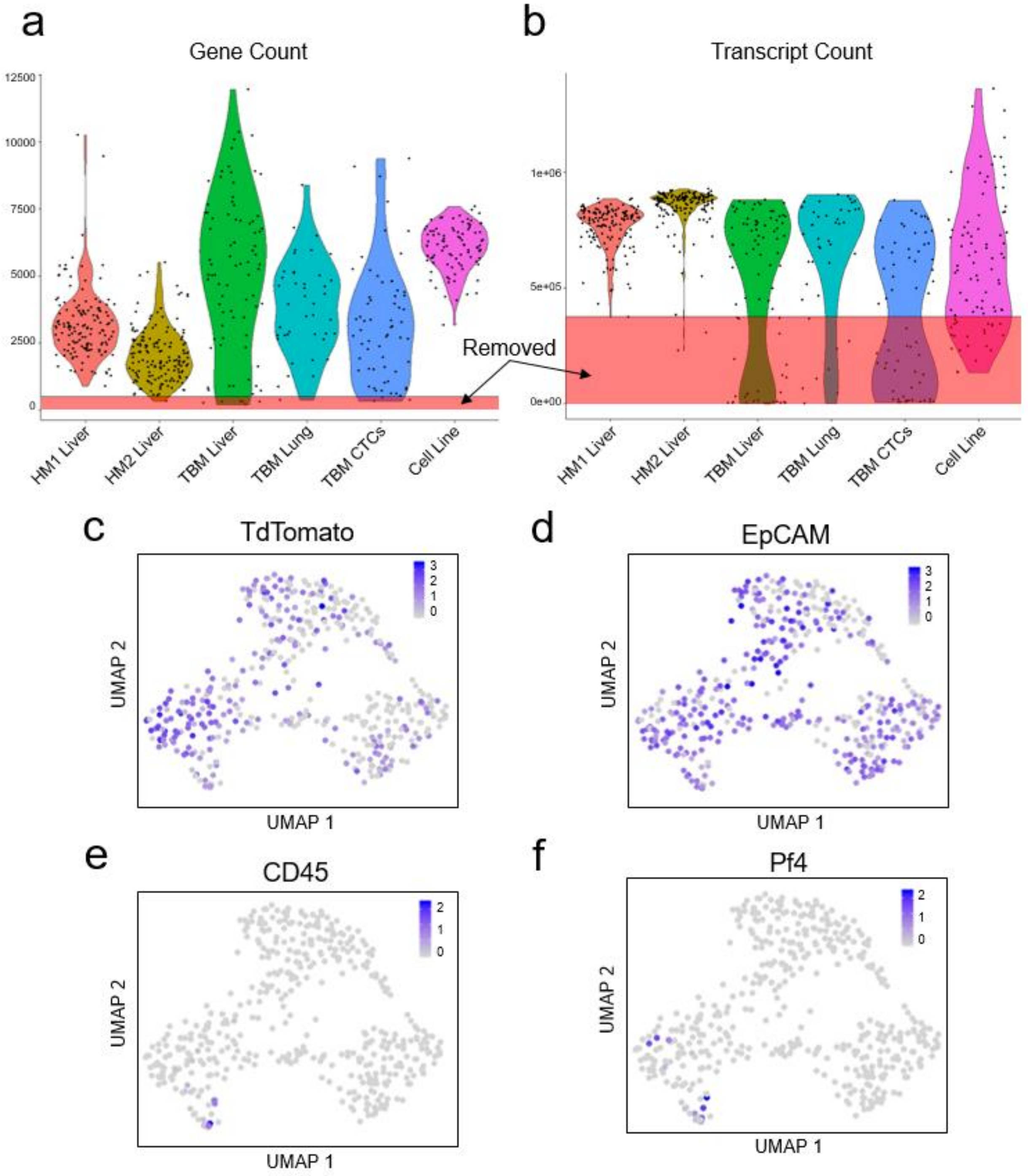


**Supplementary Fig. 8 |** Quality control and pre-processing and verification of target tumor cell identity for downstream tumor-specific analyses. Gene count (**a**) and transcript count (**b**) for each scRNA-seq sample in the study, as well as the cutoffs used for downstream analysis (gene count > 500 and transcript count > 375,000). Pre-processing feature plot to demonstrate sequenced cells are from the fluorescent tumors and blood cells have been effectively removed out. (**c - f**) UMAP feature plots showing positive expression of expected tumor cell genes (**c**) tdTomato and (**d**) EpCAM, and no expression of white blood cell marker CD45 (**e**) or platelet marker Pf4(**f**). Scale represents log-transformed expression.

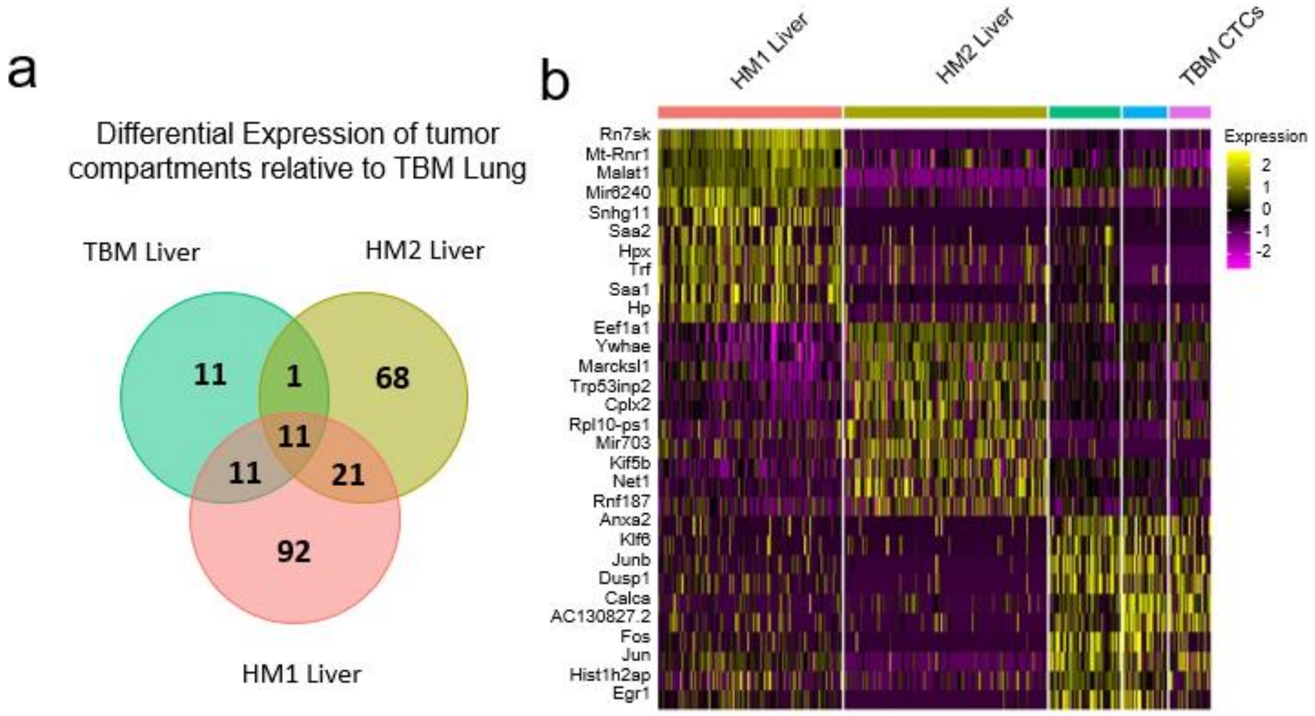


**Supplementary Fig. 9 |** (**a**) Venn diagram of genes that are differentially expressed between the TBM lung and each of the 3 liver metastases (Bonferroni adjusted $p<.05$, log2-fold change>.6) shows some variation, but also a consistent set of similarly enriched genes. (**b**) Heatmap of UMAP cluster defining genes showing inter-mouse heterogeneity. Scale bar represents scaled expression.

| | CTCs/mL | |
|---|---|---|
| Exp # | Real-Time Scan | Terminal Blood |
| 1 | 400 | 318 |
| 2 | 893 | 505 |
| 3 | 287 | 296 |
| 4 | 555 | 458 |

**Supplementary Table 1 |** Comparison between CTC concentrations as measured through real-time scans using the CTC counter and terminal blood scans collected after the real-time scan from the same animal.
